# Development of Machine Learning Model for Pan-cancer Subgroup Identification using Multi-omics Data

**DOI:** 10.1101/2022.09.15.507989

**Authors:** Seema Khadirnaikar, Sudhanshu Shukla, S R Mahadeva Prasanna

## Abstract

Cancer is a heterogeneous disease and patients with tumors from different organs can share similar epigenetic and genetic alterations. Therefore, it is crucial to identify the novel subgroup of patients with similar molecular characteristics. It is possible to propose a better treatment strategy when the heterogeneity of the patient is accounted for during subgroup identification irrespective of the tissue of origin. In this work, mRNA, miRNA, DNA methylation, and protein expression features from pan-cancer samples were concatenated and non-linearly projected to lower dimension using machine learning (ML) algorithm. This data was then clustered to identify multi-omics based novel subgroups. The clinical characterization of these ML subgroups indicated significant differences in overall survival (OS) and disease free survival (DFS) (p-value*<*0.0001). The subgroups formed by the patients from different tumors shared similar molecular alterations in terms of immune microenvironment, mutation profile, and enriched pathways. Further, decision-level and feature-level fused classification models were built to identify the novel subgroups for unseen samples. Additionally, the classification models were used to obtain the class labels for the validation samples and the molecular characteristics were verified. To summarize, this work identified novel ML subgroups using multi-omics data and showed that patients with different tumor types can be similar molecularly. We also proposed and validated the classification models for subgroup identification. The proposed classification models can be used to identify the novel multi-omics subgroups and the molecular characteristics of each subgroup can be used to design appropriate treatment regimen.

## 1 Introduction

The recent advances in therapy protocol has increased the five-year prognosis of many cancer types, yet, cancer remains the second most common cause of death globally (Giaquinto et al., 2022). Classification of cancer is crucial for the primary diagnosis of the disease. Generally, cancers are grouped together based on their organ-of-origin. Clinically, majority of the tumors with the same histological grade and organ-of-origin are treated using a similar approach (Carbone, 2020). Despite the tumors originating from the same organ and having the matching histopathological grade, patients respond differently to the therapy resulting in different survival outcomes (Marusyk et al., 2020). One of the main reasons for this differential response is the variation in the underlying genetic and epigenetic aberrations that contribute to the heterogeneity (Hirata and Sahai, 2017; Dagogo-Jack and Shaw, 2018). A better treatment regimen can be proposed if the treatment strategies account for this heterogeneity as well (Fisher et al., 2013). Hence, there is a need to identify the novel subgroups based on the genomic and epigenomic aberrations beyond the organ-of-origin. The advancement in next generation sequencing technologies have contributed to the generation of massive amount of data representing various genomic and epigenomic changes. This laid the foundation for the establishment of The Cancer Genome Atlas (TCGA), a multiplatform cancer database housing information of more than 11,000 samples from 33 cancer types. This database instigates us to understand the association of various molecular features with the phenotype. As each molecular level of evidence (omic level or datatype) provides a different set of biomarkers and insights about the aberrations, integrative analysis of multiple datatypes accounting for the non-linear interactions will result in the identification of better subgroups (Olivier et al., 2019; Ahmed, 2020). The idea of accounting for individuals’ heterogeneity for the identification of novel subgroups falls in the realm of precision therapy, which intends to use the precise knowledge of the variation in a smaller population’s genome to recommend the therapies targeting the mechanisms specific to that sub-population (Council et al., 2011).

Several works have attempted to identify the molecular subtypes within the samples from the same tumor type and have shown the presence of multiple subtypes (Baek and Lee, 2020; Chaudhary et al., 2018; Chen et al., 2017). Identification of alterations specific to the subgroups in various studies have resulted in the development of targeted therapies (Lee et al., 2018; Oh and Bang, 2020).

Samples from different organs-of-origin and different grades can also be similar molecularly. The first attempt to identify the subgroups in pan-cancer data was made by Hoadley et al., using the data from 12 tumor types to identify 11 subgroups (Hoadley et al., 2014). Through their multi-platform analysis authors showed that the samples from different organs-of-origin did cluster together. This study was based on Cluster-Of-Cluster-Assignments (COCA). COCA is a late integration technique where there is no provision to account for the interaction among the different datatypes. Recently, another study was carried out by the same group using data from 33 different tumor types (Hoadley et al., 2018). Here, the authors used iCluster to model the interactions within the datatypes and identified 28 subgroups. However, various studies have pointed out the computational complexity associated with this statistical technique to be quite high (Sathyanarayanan et al., 2020). González-Reymúndez et al., proposed a statistical technique based on penalized matrix factorization to identify eight clusters using pan-cancer data (n = 5,408) (González-Reymúndez and Vázquez, 2020). Here, the authors have used sparse singular value decomposition (sSVD) to reduce the dimension of data and density-based spatial clustering of applications with noise (DBSCAN) algorithm to cluster the samples. As the dimension of data increases, computation of SVD gets computationally intensive.

All these pan-cancer studies are based on statistical methods. Statistical models can be used to understand the relation between the variables and draw the inferences. However, they cannot be used to identify and generalize the hidden patterns particularly in the multi-modal heterogeneous data (Ij, 2018). As the advancement in the cost-effective sequencing technologies has enabled the generation of the large dimension data at all the levels of the genome, a comprehensive analysis of this data using various machine learning (ML) techniques will help in the identification of the hidden patterns. Hence, this work aims to apply ML based techniques to multi-omics (multiple datatypes) pan-cancer data for better understanding and subgrouping of the cancer patients. Identification of molecularly similar subgroups in pan-cancer samples will help in designing better therapeutic strategies independent of the organ-of-origin.

In this work, different datatypes including mRNA, miRNA, methylation, and protein expression were integrated. To model the non-linear interaction among the molecular features from multiple datatypes, an autoencoder (AE) was trained, and a reduced dimension representation was obtained. Consensus *K*-means clustering was then carried out using this representation to identify the ML based molecular subgroups. Pairwise statistical tests were carried out in the identified subgroups using all the features. And the features specific to each subgroup were identified by filtering them based on the q-value and fold-change (FC). The chosen features were then used to train the classification models to identify the multi-omics based molecular subgroup for a new sample. Three widely used ML based classification models, support vector machines (SVM), random forest (RF), and feed-forward neural network (FFNN) were trained and their predicton probabilities were combined to obtain the decision-level fused classification models as the decision-level fused models are more robust and stable to outliers. Also, as each datatype is known to convey complementary information, we trained the classification models by combining the features from different data levels to obtain feature-level fused classification models.

## 2 Methods

The steps followed in this work for the identification of novel ML based subgroups are outlined in the Fig. 1 and explained in detail in the subsequent sections.

**Figure 1:**
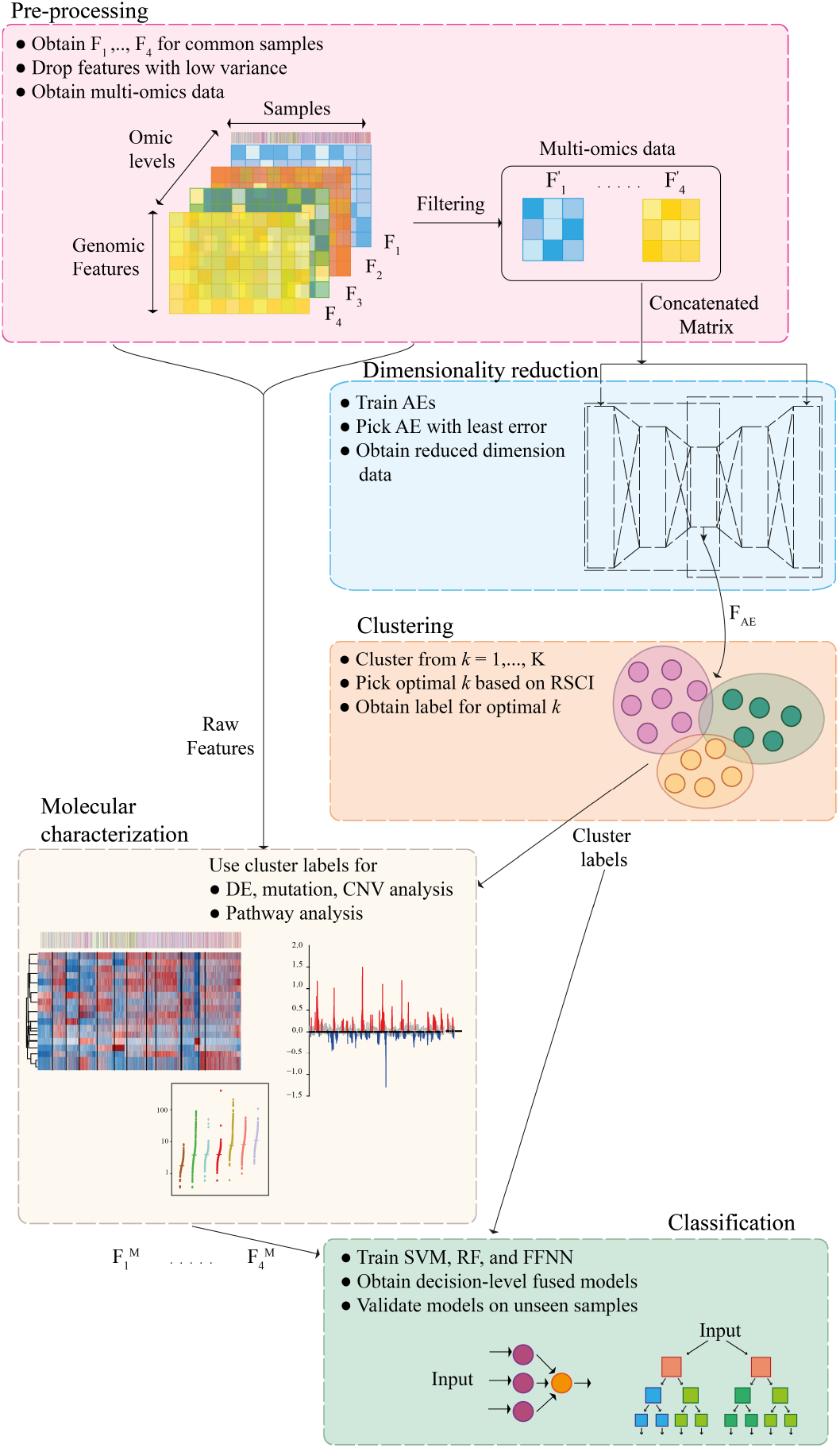
Overall pipeline followed in this work. **a)** Each datatype (single-omic) was preprocessed and multi-omics representation was obtained by stacking the features for the samples common across all the omic levels. **b)** The latent representation of multi-omics data (*F*_*AE*_) was obtained using an autoencoder (AE). **c)** Consensus *K*-means clustering was applied on the reduced dimension representation to obtain the cluster labels. **d)** To understand the subgroups obtained molecular characterization of samples in each cluster was carried out. **e)** Decision-level fused classifiers were obtained by combination of classification models including, support vector machines (SVM), random forest (RF), and feed-forward neural network (FFNN) for multi-omics subgroup identification.

### 2.1 Datasets and data preprocessing

Pan-cancer study is a collection of tumor and normal data from 33 different tumor types for multiple datatypes. We obtained RNA-sequencing FPKM values, miRNA RPKM values, and DNA methylation 450k beta values, and protein expression values (RPPA data) of solid tumors (Colaprico et al., 2016; Li et al., 2013, 2017). The abbreviation of each cancer type along with the number of samples having the information for every datatype is summarized in Supplementary Table S1. The samples (n = 5703) which had information from all the datatypes were considered for further analysis. In this work, the different datatypes are labeled as factors, and the mapping is tabulated in Supplementary Table S2. For a machine learning (ML) algorithm to learn effectively and to obtain improved generalizability, the number of samples (*n*) should be approximately equal to the number of dimensions (*p*) (i.e., *n*∼ *p*) (Bishop and Nasrabadi, 2006; Hastie et al., 2009). To satisfy this condition, preprocessing was carried out as outlined in the Supplementary Fig. S1, following the protocols from the previous studies (Chen et al., 2017; Jiang et al., 2020; Capper et al., 2018; Maros et al., 2020; Maksimovic et al., 2016). Briefly, dimensions in *F*_1_ and *F*_2_ with zeros in more than 20% of the samples were dropped (Baek and Lee, 2020). Dimensions in *F*_1_ were then sorted based on the standard deviation in the decreasing order and the top 2000 most variable dimensions were considered for further analysis. In the case of *F*_3_, dimensions (probes) with data missing in more than 10% of the samples and those on X and Y chromosomes were dropped. And similar to *F*_1_, the dimensions were sorted based on the standard deviation, and the 2000 most varying probes were considered for further analysis (Peters et al., 2015). For *F*_4_, proteins with data missing in more than 10% of the samples were dropped. In the case of both *F*_3_ and *F*_4_, NAs were imputed by K-nearest neighbors (KNN) (*K* = 5) as in previous studies (Chaudhary et al., 2018; Chen et al., 2017; Jiang et al., 2020). The selected dimensions were then stacked to obtain the multi-omics data for samples common across all the datatypes. To better understand each subtype, mutation data (maf files) and copy number data (segment data) were downloaded from TCGA. Further details including stromal fraction, and leukocyte fraction were obtained from the previously published pan-cancer studies (Hoadley et al., 2018; Thorsson et al., 2018).

### 2.2 Multi-omics data integration and cluster identification

The multi-omics data obtained by concatenating different datatypes was further reduced to a lower dimension using autoencoder (AE) to capture the non-linear interaction among the different dimensions (Pavlidis et al., 2002; Cantini et al., 2021; Ashworth et al., 2011). The number of nodes in the hidden layers and the bottleneck layer were varied to obtain multiple architectures of AE which were trained with different learning rates (Supplementary Table S3). The AE models were trained with Adam optimizer, and mean-squared error as a loss with early stopping criteria, i.e., model training was stopped if there was no reduction in validation error for ten subsequent epochs. The reduced dimension data from AE was then used to identify the novel subgroups by clustering.

Consensus *K*-means clustering was applied to the reduced dimension data. The number of clusters (*K*) was varied from 2 to 20. Clustering was repeated 1000 times with 80% of the samples chosen randomly (Monti et al., 2003). The proportion of samples those cluster together during different runs indicate the consistency of clustering. Proportion of ambiguously clustered pairs (PAC), the value quantified with the aid of the cumulative distribution function (CDF) curve is one such variable (Şenbabaoğlu et al., 2014). The section lying in between the two extremes of the CDF (*C*) curve (*u*_1_ and *u*_2_, Supplementary Fig. S3 (a)) quantifies the proportion of samples that were assigned to different clusters in each iteration. PAC (*P*) is used to estimate the value of this section representing the ambiguous assignments and is defined by (1), where *K* is the desired number of clusters.

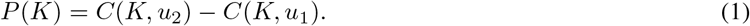

Lower the value of PAC, lower is the disagreement in clustering during different iterations or in other words, more stable are the clusters obtained (Şenbabaoğlu et al., 2014). As PAC does not account for a reference distribution, it is biased towards the higher values of *K*. Relative cluster stability index (RCSI) overcomes this by generating a reference model following the Gaussian distribution without actual clusters (i.e., *K* = 1). The reference data is generated using Monte-Carlo simulation maintaining the correlation between the dimensions (John et al., 2020). Hence, RCSI was used in this work to find the optimal number of clusters. For a given number of clusters *K*, RCSI is given by (2).

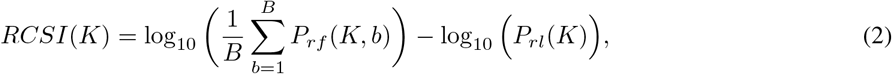

where *B* is the total number of Monte Carlo simulations, *P*_*rf*_ (*K, b*) is the PAC value of the reference distribution for *b*^th^ Monte Carlo simulation, and *P*_*rl*_(*K*) is the PAC value for the actual data. Higher the value of RCSI, the better the clustering. In this work, the values of *u*_1_ and *u*_2_ were set to 0.1 and 0.9, respectively as suggested by the authors (John et al., 2020).

### 2.3 Molecular characterization of subgroups

To check the phenotypic differences in the ML subgroups identified, logrank test was carried out using overall survival (OS) and disease-free survival (DFS) as end points and the survival difference was visualized using the Kaplan-Meier (KM) curves. Further, to understand and identify the features that define and are specific to each ML subgroup, ANOVA and pairwise t-test were used. The steps followed while preprocessing and testing each datatype are outlined in Supplementary Fig. S4. The features with *log*_2_(*FoldChange*) ≥5 and *q* ≤0.01 were considered to be differentially expressed and used for further interpretation. To identify the gene ontology (GO) pathways enriched in each ML cluster, Metascape analysis was carried out using the protein-coding genes (PcGs). Also, the expression of the PcGs were used to carryout the Gene Set Enrichment Analysis (GSEA) using the hallmark geneset to better understand the various pathways enriched in each subgroup (Subramanian et al., 2005; Mootha et al., 2003). The leukocyte fraction, and stromal fraction for each sample obtained from the previous studies were also used to gain further insights about the subgroups (Thorsson et al., 2018; Hoadley et al., 2018). To understand the tumor microenvironment (TME), CIBERSORT analysis was carried out using the LM22 signature gene set (Newman et al., 2015). To obtain the mutation signatures the steps described by the previous studies were followed (Hoadley et al., 2018; Covington et al., 2016). To summarize, the mutation signature for each sample was obtained by deconvolution of the mutation matrix with the signature weight matrix by non-negative matrix factorization (NMF) (Covington et al., 2016). The weight matrix representing each mutation signature was obtained from the previous study (Covington et al., 2016). To understand the copy number variation, G-score which accounts for the amplitude of copy number variation and the frequency of occurrence of aberration in each cytoband was obtained from GISTIC analysis and was visualized using Maftools (Mermel et al., 2011; Mayakonda et al., 2018). To deduce the pathway activity associated with each ML subgroup, details of 22 gene programs and 20 drug program were obtained from previous studies and analyzed by plotting their average values (Hoadley et al., 2018, 2014).

### 2.4 Subgroup identification by classifier combination

The features specific to each subgroup identified by the molecular analysis in the previous section along with the subgroup labels obtained by clustering were used to train the ML models to identify the subgroups for a new sample. Three ML models, support vector machine (SVM), random forest (RF), and feed-forward neural network (FFNN) were trained. The hyper-parameters were tuned by five-fold cross-validation repeated ten times. SVM was trained with a radial kernel as the interaction among the different dimensions is known to be non-linear. FFNN was trained with different learning rates (0.001, 1e-04, and 1e-05), and the optimal one was chosen by a five-fold CV. Train-test split of 90%−10% was used to build all the models. Besides the individual classifiers (*L*_0_), decision-level fused classifiers (*L*_1_) were also built to obtain precise predictions. To train these models, the prediction probabilities obtained from individual classifiers (*P*_*SV M*_, *P*_*RF*_, and *P*_*FFNN*_) were stacked and used as input, and the cluster labels as output. A linear decision-level fused model was obtained by linearly weighing the prediction probabilities from individual *L*_0_ classifiers by α, *β*, and *γ*, respectively (Pavlidis et al., 2002; Potamianos et al., 2003; Oh and Kang, 2017). The final prediction (*P*_*F*_) was obtained by the weighted summation of individual prediction probabilities using (3) (Rabha et al., 2019).

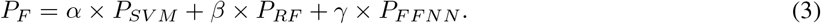

The values of α, *β*, and *γ* were varied from 0 to 1 in steps of 0.01 by ensuring that they sum up to 1 (Algorithm S1).

Rather than manually iterating through a set of weights to identify the significance of each classifier, ML models like logistic regression and FFNN were also trained using the prediction probabilities from individual classifiers to identify the multi-omics subgroups. As non-linear models were used to combine the prediction probabilities, we called the resulting models as non-linear decision-level fused models. If the same set of samples are used to train both *L*_0_ and *L*_1_ classifiers, the performance of the classifier will degrade when the distribution of test set is different from the training set. To overcome this, in this work, two sets of decision-level fused models were trained.

## 3 Results

Datatypes including mRNA, miRNA, DNA methylation, and protein expression were integrated and subgroups were identified following the protocol outlined in Figure 1. Number of samples and dimensions considered for analysis from various tumor types are tabulated in Supplementary Table S1 and S2.

### 3.1 Dimensionality reduction and clustering of pan-cancer data

The multi-omics representation of pan-cancer samples was obtained by concatenation of data from different datatypes (Supplementary Fig. S1). In a complex disease like cancer, there often exists a non-linear interaction among the different molecular features (Hira and Gillies, 2015; Alanis-Lobato et al., 2015). To capture this non-linear interaction, multiple autoencoder (AE) models were trained and the model with the least difference in the training and validation loss was chosen to reduce overfitting (Supplementary Table S3) (Goodfellow et al., 2016). The reduced dimension data from the AE model was then clustered by consensus *K*-means clustering to identify the subgroups across the TCGA tumors. RCSI value, which takes a null reference distribution to calculate PAC was used to pick the optimum number of clusters as described in the methods section. The cluster with the highest RCSI value (*K* = 13) was considered for further analysis (Fig. 2 (a) and (b), Supplementary Fig. S2 (a)). To visualize the distribution of samples, tSNE plots were plotted using the reduced dimension AE data. The samples in the tSNE plots were colored based on the tumor type and the cluster numbers (C1 to C13) (Fig. 2 (c) and (d)). The tSNE plot labelled with 13 ML clusters as identified in this analysis were more compact and well separated (silhouette width = 0.43) than the clusters obtained based on the tumor type (silhouette width = 0.32), showing the superiority of the stratification when carried out using the proposed method.

**Figure 2:**
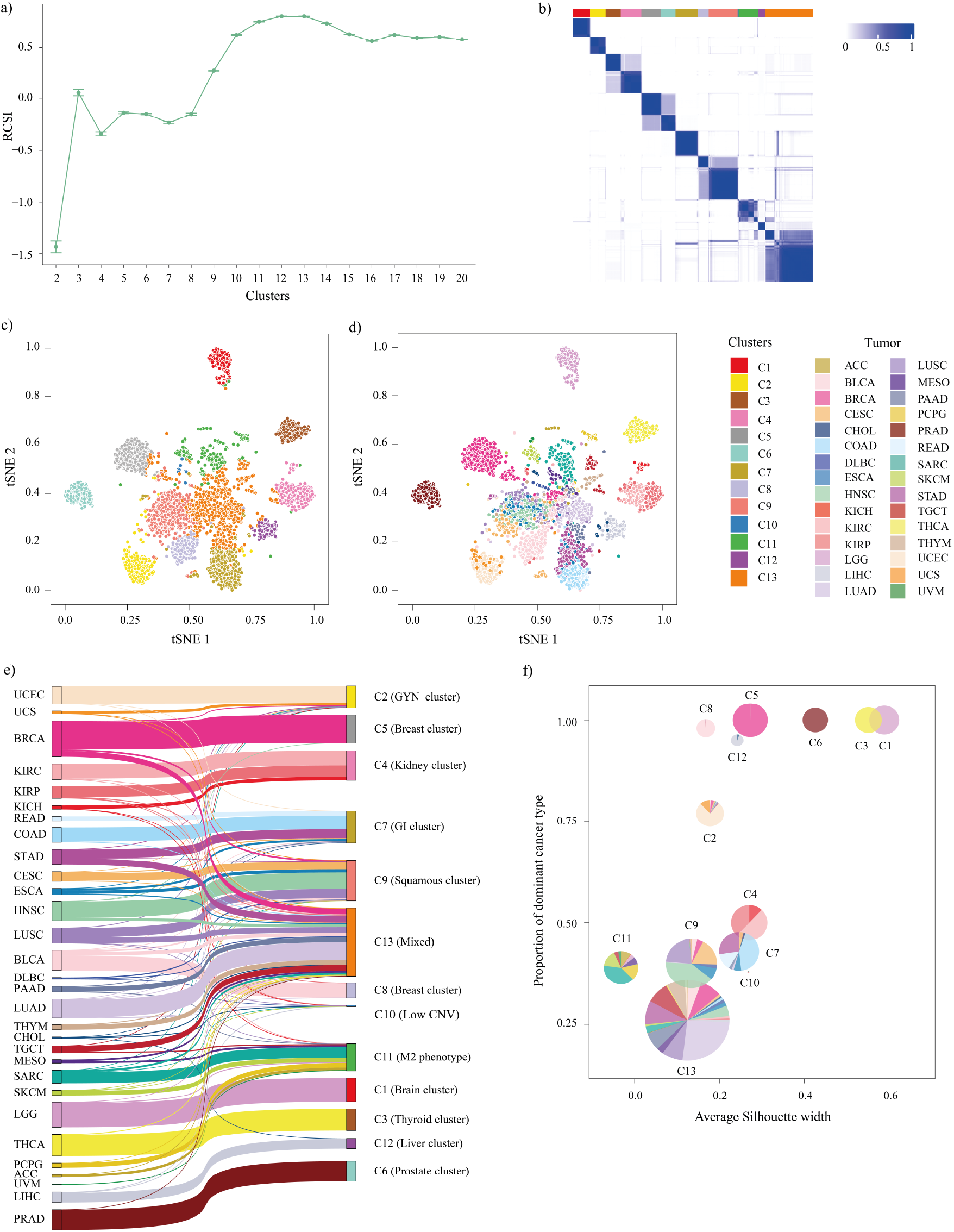
Composition and distribution of samples in ML clusters. **a)** RCSI values for *K* = 2 to *K* = 20. **b)** Consensus heatmap plot for consensus *K*-means clustering (*K* = 13). **c) and d)** t-SNE plots for reduced dimension obtained using AE. Samples are colored based on the labels obtained by consensus *K*-means clustering and tumor type, respectively. **e)** Sankey plot depicting the relationship of samples in each cluster with TCGA tumor type. **f)** Pie chart showing the proportion of samples from various tumor types in each cluster plotted against the average silhouette width showing the compactness of each cluster.

To validate that dimensionality reduction is essential and the interaction between the molecular features is indeed non-linear, the dimension of the selected raw features was reduced by the most widely used linear dimensionality reduction technique, PCA. And the features representing 99% of the variance were retained and used for clustering. The RCSI values obtained for dimensionality reduction by PCA (RCSI = 0.14) was smaller than that obtained by AE (RCSI = 0.8). This indicated that the clusters resulting from the dimensionality reduction by AE were better and more consistent. To confirm that dimensionality reduction is crucial and the clusters obtained using multiple data types are indeed more consistent than clusters obtained using single data type, we repeated the whole pipeline for each data type using the selected features with and without dimensionality reduction. The resulting RCSI values indicated that dimensionality reduction an essential step and the clusters accounting for multiple data types were more stable than single data type (Table 1).

**Table 1:** Summarizing the RCSI values obtained for *K* = 13 for each level of evidence for the subset of selected features, when clustered without dimensionality reduction, and with dimensionality reduction using PCA and AE (*F*_1_: mRNA expression, *F*_2_: miRNA expression, *F*_3_: DNA methylation, *F*_4_: protein expression).

| Omic | Dimension | Without<br>dimensionality<br>reduction | With<br>dimensionality<br>reduction |  |
| --- | --- | --- | --- | --- |
|  |  |  | PCA | AE |
| $F_1$ | 2000 | -0.69 | -0.86 | -0.19 |
| $F_2$ | 1753 | 0.19 | 0.23 | 0.25 |
| $F_3$ | 2000 | 0.35 | -0.45 | 0.43 |
| $F_4$ | 216 | 0.48 | -0.34 | 0.31 |
| $F_1 + F_2 + F_3 + F_4$ | 5969 | 0.27 | 0.14 | 0.8 |

### 3.2 Composition of pan-cancer clusters

To further understand the composition of the ML clusters, we calculated the proportion of samples from different tumor types in each cluster (Supplementary Fig. S3). Among the 13 ML clusters, six clusters were pure with all the samples from the same tumor type, and the rest were mixed clusters with the samples from different tumor types. The ML cluster C1 was purely composed of LGG samples. Similarly, the ML clusters C3, C5, C6, C8, and C12 comprised of samples from THCA, BRCA, PRAD, BLCA, and LIHC, respectively. With samples from 27 different tumor types, ML cluster C13 was the most heterogenous and also the largest subgroup. ML cluster C10 was the smallest heterogenous cluster with the majority of the samples from KIRP, CHOL, and LIHC. ML cluster C2 had the samples from uterine, breast, and cervical cancers. Hence, we called it GYN cluster. We named ML cluster C4 as kidney cluster as it was formed by the samples from various kidney tumors. COAD, STAD, and READ accounted for the majority of the samples in ML cluster C7 and hence, named as gastrointestinal cluster (GI cluster) cluster. Interestingly, samples from HNSC, LUSC, and CESC constituted the majority of samples in ML cluster C9 which was called squamous cluster. ML cluster C11 was a mixed cluster formed by samples from 12 tumor types, with the majority of them from SARC, SKCM, and PCPG. This distribution can be seen in the Sankey plot as well which depicts the relation between the samples from different clusters and their tumor type (Fig. 2 (e)).

To understand the compactness and homogeneity within each ML cluster, the proportion of samples from the dominant tumor type in each cluster was plotted against the average silhouette width of each cluster (Fig. 2 (f)). Here, the radius of each pie is proportional to the number of samples in each cluster. Though the ML clusters dominated by a single tumor type C1 (Brain cluster), C3 (Thyroid cluster), C6 (Prostate cluster) had the highest silhouette width, ML clusters C4 (Kidney cluster) and C7 (GI cluster), which are mixed clusters, had silhouette widths closer to that of ML cluster C5 (Breast cluster), which is formed by a single tumor type. This result strengthens the hypothesis that, despite originating from different tumor types, the samples in these ML clusters are close to each other molecularly.

### 3.3 Clinical and biological characterization of clusters

To analyze if there exists any differences in the survival times (overall survial (OS) and disease-free survial (DFS)) between the 13 ML clusters obtained, log-rank test carried out using the was used to compare the survival times and Kaplan-Meier (KM) plots to visualize the survival curves. This analysis indicated that there exists at least one group with significantly different survival time when compared to the others (OS and DFS p-value < 0.0001, Supplementary Fig. S2 (b) and (c)). With an aim to gain further insights into the distinguishing features of each ML cluster contributing for the variation in phenotype, statistical tests were carried out as described in methods section. This analysis identified 2868 protein-coding genes (PcGs), 442 long non-coding RNAs (lnc RNAs), 104 miRNAs, 4872 methylation probes, and 216 proteins to be differentially expressed (DE) (Fig. 3 and Supplementary Table S5). To interpret the function associated with various genes identified, PcGs identified within each ML cluster were used to carryout the metascape analysis and the top 5 GO biological processes were tabulated (Supplementary Table S6). ML clusters C5 (Breast cluster), C4 (Kidney cluster), and C7 (GI cluster) had activation of pathways associated with embryonic morphogenesis, circulatory system process, and epithelial cell differentiation, respectively. Further, GSEA analysis was carried out using the hallmark genesets to identify the genesets enriched in each ML cluster. As expected, the pathways associated with estrogen and androgen response were positively enriched in ML clusters C5 (Breast cluster) and C6 (Prostate cluster), respectively (Supplementary Table S7, and Supplementary Table S8).

**Figure 3:**
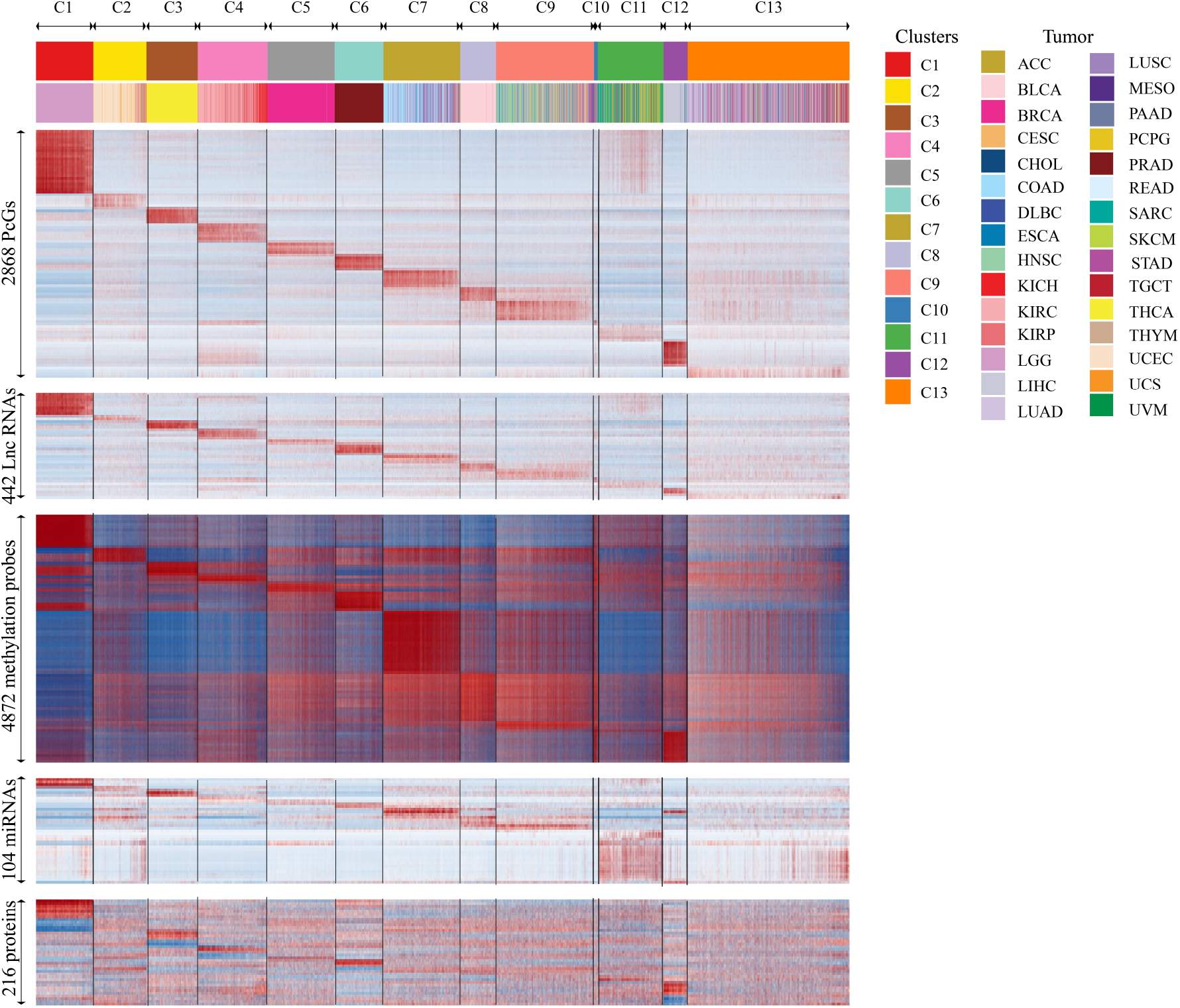
Characterization of different molecular levels of evidence in ML clusters. Heatmap indicating the expression of protein coding genes (PcGs), long non-coding RNAs (Lnc RNAs), methylation probes, miRNAs, and protein expression in the subgroups obtained by multi-omics clustering.

Immunotherapy has become a crucial treatment option for many tumor types (Esfahani et al., 2020; Waldman et al., 2020). The response to immunotherapy partially depends on the immune microenvironment of the tumor (Murciano-Goroff et al., 2020; Tang et al., 2021; Petitprez et al., 2020). Hence, we analyzed the proportion of stromal (non-tumor) and leukocyte cells in each ML cluster (Fig. 4). ML cluster C13 (Mixed) showed the highest infiltration of stromal and leukocyte cells (Fig. 4 (a) and (b)). To understand the proportion of leukocytes contributing to the stromal fraction, the leukocyte fraction was plotted against the stromal fraction (Fig. 4 (c)). In case of the samples those are close to the diagonal or along the diagonal, leukocytes contribute for the majority of the stromal proportion (Hoadley et al., 2018). Hence, C13 had an immune rich microenvironment where the majority of the stromal fraction was contributed by the leukocytes. To understand the difference in immune microenvironment of each ML cluster, Cibersort analysis was performed using the LM22 geneset. We found that many clusters showed unique inflitration of various immune cells (Supplementary Fig. S5 and Supplementary Table S9). ML cluster C13 was observed to have a high infiltration of naive CD4 T cells which have a vital role play in immune response (Luckheeram et al., 2012; Caccamo et al., 2018). ML cluster C11 had the highest infiltration of M2 macrophages and hence was named M2 phenotype.

**Figure 4:**
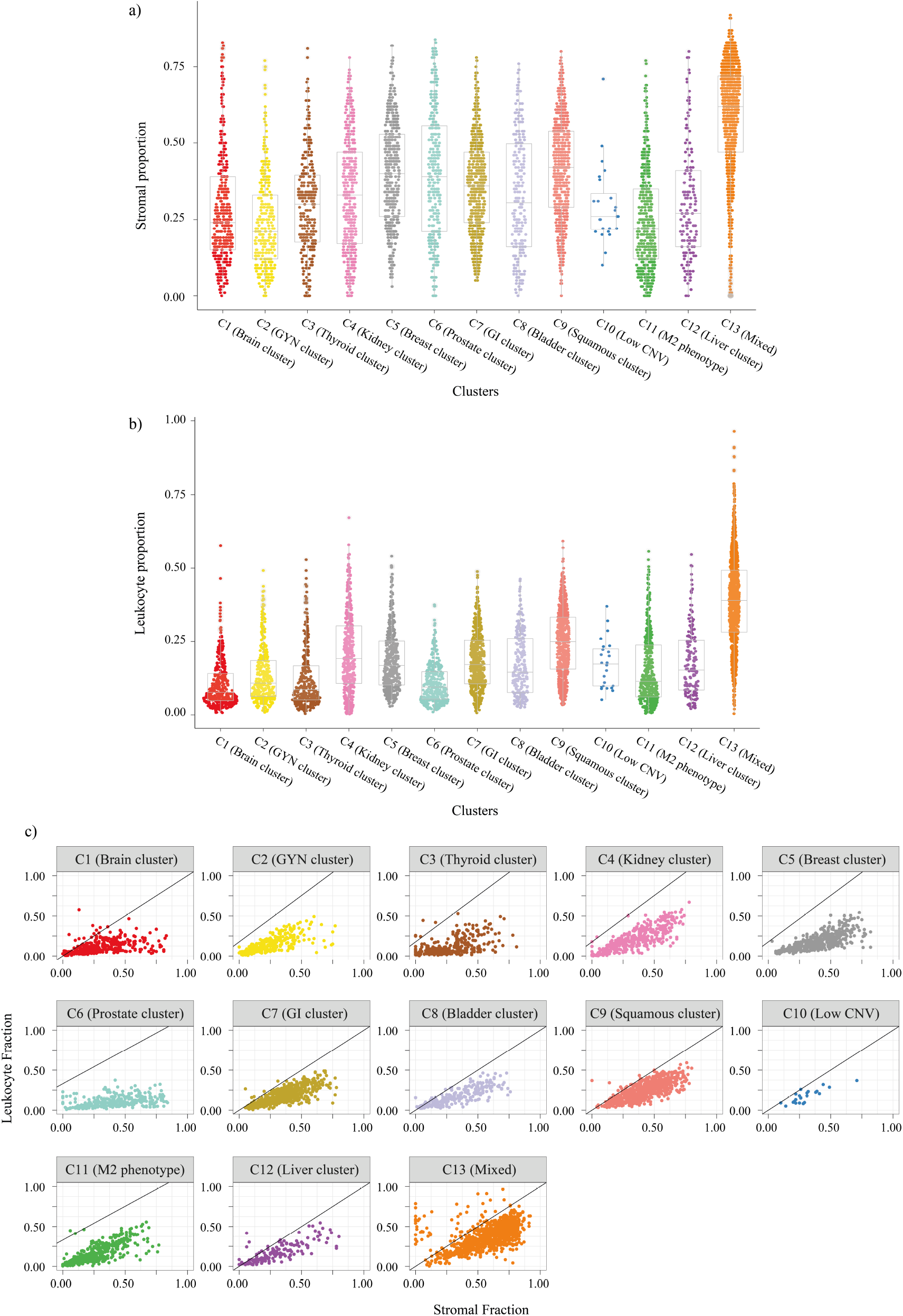
Quantifying the tumor microenvironment of ML clusters. **a)** Stromal fraction for pan-cancer samples in each cluster. **b)** Leukocyte fraction for pan-cancer samples in each cluster. **c)** Leukocyte vs. Stromal fraction for pan-cancer samples in each cluster.

As mutations have a significant role in the tumor development and progression, non-silent mutation rates, and mutation signatures were obtained for all the ML clusters (Fig. 5). We found a significant variation in mutation rates of each cluster. Interestingly, we found that ML cluster C8 (Bladder cluster) showed the highest mutation burden and ML cluster C3 (Thyroid cluster) had the lowest median non-silent mutation rate (Fig. 5 (a)). The mutation signature obtained by non-negative matrix factorization (NMF) as described in methods section showed the different characteristics associated with each ML cluster (Fig. 5 (b)). Signature 6 was the signature most commonly expressed across all the samples. It represents the CpG mutations indicating high enrichment of C*>*T mutations. ML cluster C2 (GYN cluster) had enrichment of MutSig 19 (POLE) indicating the activity of polymerase-E which is generally associated with the endometrial and colorectal cancers (Xing et al., 2022). C2 was composed of UCEC, a subtype of endometrial carcinoma known to have POLE mutations. GI cluster (C7) was enriched for mutation signature associated with arsenic activity. APOBEC family of enzymes are known to cause mutation in various cancers (Petljak and Maciejowski, 2020). We found enrichment of APOBEC mutation signature in Bladder cluster (C8) and squamous cluster (C9) (Fig. 5 (b)). We also found enrichment of UVB mutation signature in ML cluster C11 which consists of SKCM and SARC cancers. ML cluster C13 (Mixed) with the majority of samples from LUAD (26%) had enrichment of mutation signature associated with smoking. To understand the copy number variation in each ML cluster, the G-score obtained by GISTIC analysis was visualized using Maftools. We identified the top five most frequently aberrated cytobands in each cluster and plotted (Supplementary Fig. S6). The analysis showed high CNV in ML cluster C2 (GYN cluster), C5 (Breast cluster), C7 (GI cluster), and C8 (Bladder cluster) and very minimal CNV in C3 (Thyroid cluster), C4 (Kidney cluster), C6 (Prostate cluster), and C10 (Low CNV).

**Figure 5:**
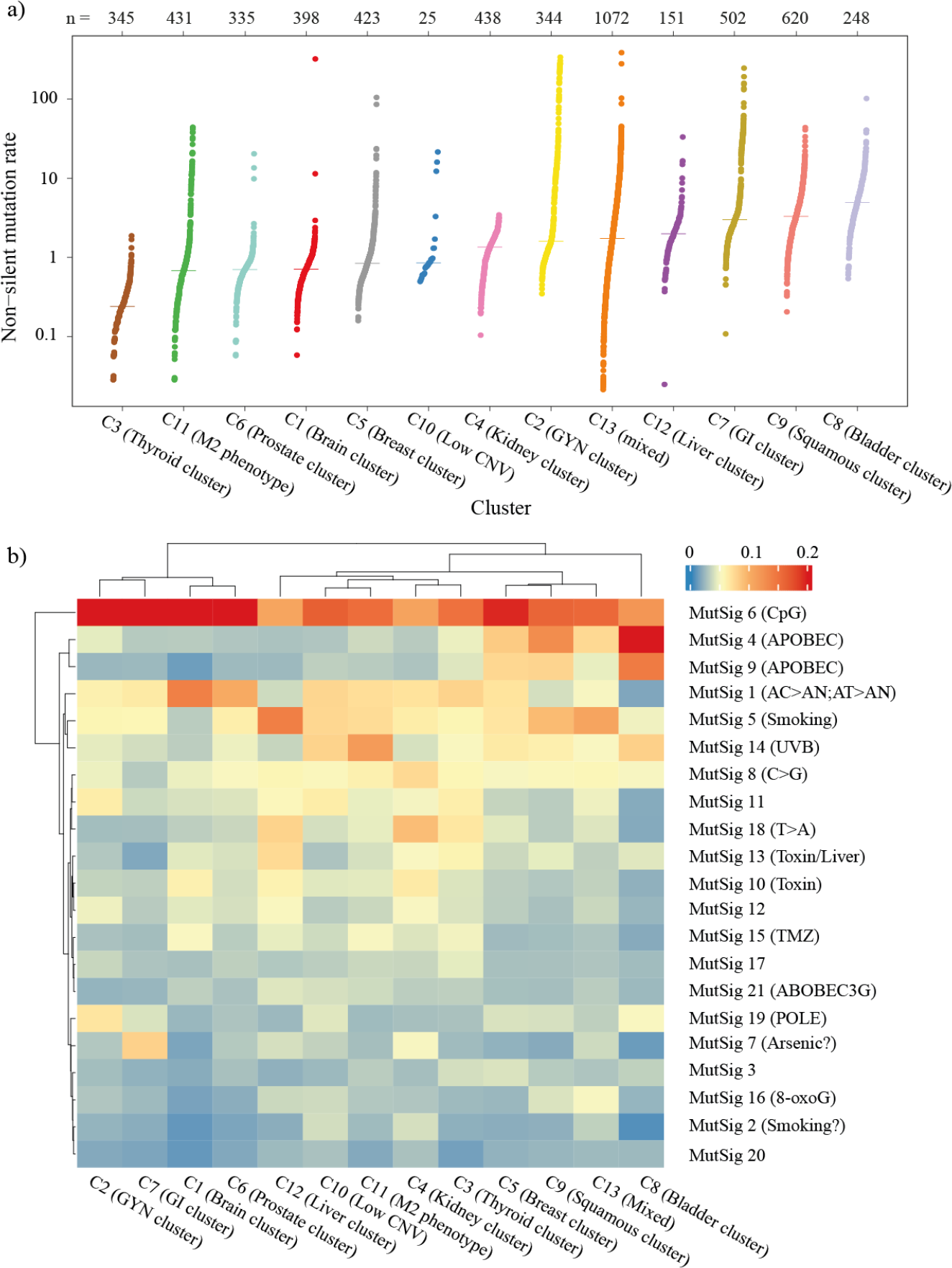
Mutation analysis of ML clusters. **a)** Non-silent mutation in each cluster sorted by median. **b)** Heatmap showing average value of mutational signature in each cluster.

Further, to interpret the pathway activity, details of the genes associated with 22 gene programs and 20 drug programs were obtained from previous work and their average value was plotted (Supplementary Fig. S7) (Hoadley et al., 2018). ML cluster C1 (Brain cluster) had the activity of gene programs associated with FOXO stemness, Neural signaling, and tumor suppressing. ML clusters C2 (GYN cluster) and C7 (GI cluster) had higher expression of genes associated with cell-cell adhesion MClaudin cluster, C4 (Kidney cluster) of hypoxia glycolysis, C8 (Bladder cluster) of 1Q amplicon, and C13 (Mixed) of Immune T-cell B-cell. ML cluster C3 (Brain cluster) showed activation of the drug pathways associated with the ALK pathway and MYC amplified chr8q24. ML cluster C6 (Prostate cluster) had the activity of Nelson response to androgen up, and C12 (Liver cluster) of KEGG Retinol metabolism. Genes associated with Heller HDAC targets were expressed in ML clusters C8 (Bladder cluster) and C9 (Squamous cluster), and CTLA4 and PD1 signaling in ML clusters C4 (Kidney cluster) and C13 (Mixed).

### 3.4 Subgroup identification by classifier combination

As all the studies do not have information from all the datatypes, classifiers were built using individual datatypes to identify the subgroups for a new sample that might have information from one datatype only. Here, three classifiers support vector machines (SVM), random forest (RF), and feed-forward neural network (FFNN), each based on different working principle were trained. The prediction probabilities obtained from them were combined using linear and non-linear models to obtain linear and non-linear decision-level fused models. All the decision-level fused models were obtained for train and test split with and without holdout as described in the methods section to obtain better generalized models. The accuracies obtained for different classifiers indicated the highest classification accuracy for the models built using *F*_3_ (DNA methylation) data type for both individual and decision-level fused models (Table 2 and Table 3).

**Table 2:** Summarizing the test accuracy of *L*_0_ classifiers for different levels of evidence (*F*_1_: mRNA expression, *F*_2_: miRNA expression, *F*_3_: DNA methylation, *F*_4_: protein expression, *F*_*AE*_: features from bottleneck layer of autoencoder, SVM: support vector machine, RF: random forest, FFNN: feed-forward neural network).

| Omic level | Dimension | w/o holdout |  |  | with holdout |  |  |
| --- | --- | --- | --- | --- | --- | --- | --- |
|  |  | SVM | RF | FFNN | SVM | RF | FFNN |
| $F_1$ | 3311 | 87.698 | 89.455 | 91.564 | 86.995 | 88.928 | 89.807 |
| $F_2$ | 104 | 86.292 | 76.098 | 84.183 | 84.534 | 76.45 | 80.492 |
| $F_3$ | 4872 | 93.849 | 95.431 | 96.661 | 92.794 | 94.903 | 92.97 |
| $F_4$ | 216 | 88.225 | 86.467 | 81.195 | 88.225 | 85.94 | 85.589 |
| $F_3 + F_1$ | 8183 | 80.492 | 94.2 | 95.782 | 74.692 | 94.728 | 96.485 |
| $F_3 + F_2$ | 4976 | 93.673 | 95.079 | 96.661 | 92.619 | 94.903 | 96.661 |
| $F_3 + F_4$ | 5088 | 93.322 | 94.903 | 95.606 | 92.267 | 94.025 | 97.012 |
| $F_{AE}$ | 100 | 98.594 | 97.54 | 97.891 | 97.188 | 97.012 | 95.782 |

**Table 3:** Summarizing the test accuracy of *L*_1_ classifiers for different levels of evidence (*F*_1_: mRNA expression, *F*_2_: miRNA expression, *F*_3_: DNA methylation, *F*_4_: protein expression, *F*_*AE*_: features from bottleneck layer of autoencoder, LR: logistic regression, FFNN: feed-forward neural network).

| Omic level | Dimension | w/o holdout |  |  | with holdout |  |  |
| --- | --- | --- | --- | --- | --- | --- | --- |
|  |  | Linear | LR | FFNN | Linear | LR | FFNN |
| $F_1$ | 3311 | 91.916 | 91.564 | 92.443 | 91.916 | 92.443 | 92.091 |
| $F_2$ | 104 | 76.098 | 82.074 | 75.22 | 85.413 | 86.292 | 83.656 |
| $F_3$ | 4872 | 97.715 | 97.364 | 97.364 | 96.485 | 96.661 | 96.661 |
| $F_4$ | 216 | 86.643 | 85.062 | 83.831 | 88.049 | 87.346 | 88.928 |
| $F_3 + F_1$ | 8183 | 96.309 | 94.376 | 95.606 | 96.134 | 95.958 | 95.606 |
| $F_3 + F_2$ | 4976 | 97.54 | 97.012 | 97.364 | 96.661 | 96.661 | 95.079 |
| $F_3 + F_4$ | 5088 | 96.661 | 96.661 | 96.309 | 97.715 | 97.188 | 96.485 |
| $F_{AE}$ | 100 | 98.946 | 98.77 | 98.77 | 97.891 | 97.364 | 95.431 |

As each molecular level (datatype) is known to carry and convey different and complementary information, the dimensions from each level were fused with the information from *F*_3_ (as it had the highest classification accuracy) to obtain feature-level fused models. We did not observe a significant improvement in the accuracy of feature-level fused classification models as compared to individual classification models (Table 2 and Table 3).

To validate the classification models, the samples with methylation data which were not used to identify the multi-omics labels were chosen. The class labels for these samples were obtained using the previously trained classification models. Interestingly, the ovarian cancer (OV) samples which were never used during training were all assigned to C2 (GYN cluster) (Supplementary Fig. S8). Further, the gene and drug pathway analysis of these samples were obtained and validated (Supplementary Fig. S9). It was observed that the variation in the pathway activity of validation samples was similar to that of original samples. Hence, confirming the utility of the trained models to obtain the multi-omics labels even in the presence of single datatype.

## 4 Discussion

Despite the continuous improvements in the treatment strategies for cancer care, it still remains one of the leading causes of death worldwide. Though the existing treatment regimen has increased the overall survival time in some cancers, the five-year survival rate remains dismal for many cancer types. Current treatment strategies rely hugely on the histopathological grades which might not be most appropriate in all cases as it is difficult to visually quantify the underlying molecular variations leading to the phenotypic change. Hence, with an aim to identify the subgroups beyond the histological subtypes, we integrated and analyzed information from various datatypes using machine learning (ML) models across the pan-cancer samples. In this work, information from mRNA, miRNA, DNA methylation, and protein expression were considered for novel subgroup identification. For machine learning models to learn and recognize the patterns effectively, the condition, *n*∼ *p* (where *n* is the number of samples and *p* is the number of dimensions) must be satisfied. Hence, we used variance based filtering to ensure that the number of dimensions (*p*) is approximately equal to the number of samples (*n*).

To account for the non-linear interaction among the features, an autoencoder (AE) was trained and the data was projected to a lower-dimensional space to identify the multi-omics based molecular subgroups. Among the 13 molecular subgroups identified, six were pure i.e samples were from the same organ-of-origin, and the rest were formed by the combination of samples from different tumor types (Fig. 2 (c)). Though the ML clusters C4 (Kidney cluster) and C7 (GI cluster) were formed by samples from different tumor types, the silhouette widths of these were comparable with pure Breast cluster (C5). This analysis revealed that molecularly similar samples from different tumor types can form a cluster which is as consistent as the cluster formed by the samples from a single tumor type.

Molecular analysis of the identified subgroups indicated similarities within the ML clusters in terms of various molecular aberrations despite the samples belonging to different tumor types. ML cluster C13 (Mixed) was the most diverse cluster with samples from 27 different tumor types. It also had the highest infiltration of stromal (non-tumoral) cells, leukocyte cells, and naive CD4 T cells (Fig. 4 and Supplementary Fig. S5). These findings suggest that patients from different tissue of origin are molecularly and immunologically similar, and can be considered for similar treatment option. ML cluster C8 (Bladder cluster) had the highest median non-silent mutation rate with enrichment for the APOBEC mutation signature which is known to cause mutation in the bladder (Fig. 5). ML cluster C3 (Thyriod cluster) had the lowest median non-silent mutation rate and enrichment of genes associated with thyroid hormone generation (Fig. 5 (a) and Supplementary Table S6).

Different classification algorithms (SVM, RF, FFNN) were trained to identify the subgroup for a new incoming sample. The prediction probabilities from these models were used to build decision-level fused models as accounting for decisions from different classifiers helps in the reduction of variance in the error, making the classification model more stable and robust to outliers. The models trained on methylation data (*F*_3_) had the highest classification accuracy indicating that this data type carried the highest information required for molecular subgroup identification (Table 2 and Table 3). Classification models were also trained based on the feature-level fusion techniques to account for the interaction among the molecular features from different datatypes. These models did not show a significant improvement in the classification accuracy when compared with the single datatype models. Though the combination of features adds additional information which will help in the classification, it will also lead to an increase in the dimension of the input data. The lack of improvement in the classification accuracy for the feature-level fused models might be because the classification models were trained with the same number of samples (*n*) but with an increased number of features (*p*). The model might not be able to capture and learn the pattern as *p* ≫ *n* in the case of the feature-level fused models.

Though the classification models were built separately for each datatype, the class labels were obtained by multi-omics integrated data. We tested the proposed model on validation samples (samples with only *F*_3_ datatype) and proved that the proposed classification models can be used to identify the multi-omics molecular subgroup for a new sample even in the absence of multiple datatypes. Therefore, the subgroup identification technique proposed in this work might have clinical utility and provide additional information alongside the histological grades. Further, the molecular characteristics specific to each subgroup might help in the selection of an appropriate treatment strategy.

In this work, only a subset of features were chosen from each datatype as part of the multi-omics data which was then used for the identification of subgroups citing the limitation in terms of the number of samples. It is possible that the incorporation of additional features from the existing datatypes and also from other datatypes like whole slide histopathological images will add further information and provide more useful insights for a better understanding of cancer samples. This will be explored as part of our future work.

## 5 Conclusions

This work attempted the identification of novel molecular subgroups in pan-cancer samples using the multi-omics data. To handle the large dimensional multi-omics data, and also to capture the non-linear interactions among the various datatypes, different machine learning based techniques were applied. We identified 13 different subgroups independent of the organ-of-origin. We showed that the samples from different organs-of-origin can form the clusters which are as stable as the clusters formed by the samples from similar organ-of-origin. Molecular characterization of the subgroups thus obtained highlighted the alterations specific to each subgroup confirming the distinctness of each subgroup. An attempt was also made to build decision-level fused classification models to identify the subgroup for a new sample. Further, the classification models were validated using single datatype and the molecular characteristics of samples were verified. Taken together, we applied ML based methods on multi-omics data to identify the novel subgroups of cancer patients. Each subgroup was characterised to identify molecular changes at epigenetic and genetic levels. Also, classification models were built to classify an unseen sample in the 13 ML clusters.

## Acknowledgment

The results shown here are based upon data generated by the TCGA Research Network: https://www.cancer.gov/tcga.

