## Supplemental Table for "Development of Machine Learning Model for Pan-cancer Subgroup Identification using Multi-omics Data" (from another version of this preprint: version 1, 2022-09-17)

**Supplementary Table S1**: Table describing the TCGA tumor types and the number of samples having information from different datatypes across different tumor types.

| Study abbreviation | Study name | mRNA | miRNA | DNA methylation | Protein expression | Selected samples |
|---|---|---|---|---|---|---|
| ACC | Adrenocortical carcinoma | 79 | 80 | 80 | 46 | 45 |
| BLCA | Bladder Urothelial Carcinoma | 411 | 417 | 412 | 344 | 336 |
| BRCA | Breast invasive carcinoma | 1087 | 1096 | 782 | 874 | 605 |
| CESC | Cervical squamous cell carcinoma and endocervical adenocarcinoma | 303 | 307 | 307 | 171 | 160 |
| CHOL | Cholangiocarcinoma | 36 | 36 | 36 | 30 | 30 |
| COAD | Colon adenocarcinoma | 465 | 455 | 296 | 357 | 244 |
| DLBC | Lymphoid Neoplasm Diffuse Large B-cell Lymphoma | 48 | 47 | 48 | 33 | 33 |
| ESCA | Esophageal carcinoma | 161 | 186 | 185 | 126 | 105 |
| GBM | Glioblastoma multiforme | 156 | 0 | 140 | 205 | 0 |
| HNSC | Head and Neck squamous cell carcinoma | 500 | 523 | 528 | 346 | 323 |
| KICH | Kidney Chromophobe | 65 | 66 | 66 | 63 | 63 |
| KIRC | Kidney renal clear cell carcinoma | 534 | 544 | 319 | 445 | 259 |
| KIRP | Kidney renal papillary cell carcinoma | 288 | 291 | 275 | 208 | 203 |
| LGG | Brain Lower Grade Glioma | 511 | 512 | 516 | 427 | 418 |
| LIHC | Liver hepatocellular carcinoma | 371 | 372 | 377 | 184 | 175 |
| LUAD | Lung adenocarcinoma | 522 | 513 | 458 | 362 | 313 |
| LUSC | Lung squamous cell carcinoma | 502 | 478 | 370 | 325 | 252 |
| MESO | Mesothelioma | 86 | 87 | 87 | 61 | 60 |
| OV | Ovarian serous cystadenocarcinoma | 374 | 491 | 10 | 411 | 0 |
| PAAD | Pancreatic adenocarcinoma | 177 | 178 | 184 | 105 | 96 |
| PCPG | Pheochromocytoma and Paraganglioma | 178 | 179 | 179 | 79 | 77 |
| PRAD | Prostate adenocarcinoma | 495 | 498 | 498 | 351 | 343 |
| READ | Rectum adenocarcinoma | 166 | 161 | 98 | 130 | 75 |
| SARC | Sarcoma | 259 | 259 | 261 | 221 | 216 |
| SKCM | Skin Cutaneous Melanoma | 103 | 97 | 105 | 96 | 85 |
| STAD | Stomach adenocarcinoma | 375 | 446 | 395 | 392 | 261 |

| | | | | | | |
|---|---|---|---|---|---|---|
| TGCT | Testicular Germ Cell Tumors | 150 | 150 | 150 | 118 | 118 |
| THCA | Thyroid carcinoma | 502 | 506 | 507 | 372 | 365 |
| THYM | Thymoma | 119 | 124 | 124 | 90 | 86 |
| UCEC | Uterine Corpus Endometrial Carcinoma | 547 | 545 | 431 | 404 | 298 |
| UCS | Uterine Carcinosarcoma | 56 | 57 | 57 | 48 | 47 |
| UVM | Uveal Melanoma | 80 | 80 | 80 | 12 | 12 |
| | | 9706 | 9781 | 8361 | 7436 | 5703* |

* Samples having information from all the levels of evidence

**Supplementary Table S2:** Summarizing the datatypes, their mapping, number of samples, original dimension, and dimensions retained after preprocessing across pan-cancer samples.

| Data type | Factor | Number of samples | Original dimension | Dimensions retained |
|---|---|---|---|---|
| mRNA | $F_1$ | 9706 | 56537 | 2000 |
| miRNA | $F_2$ | 9871 | 1881 | 1753 |
| DNA Methylation | $F_3$ | 8361 | 485k | 2000 |
| Protein expression | $F_4$ | 7436 | 258 | 216 |
| | | 5704 * | 544k | 5969 |

Note: k here indicates the multiple of 1000.
* - samples common across all the levels of evidence

**Supplementary Table S3**: Training and validation losses for different architectures of AEs.

| Architecture | Training loss | Validation loss | Difference |
|---|---|---|---|
| i/p, 4000, 2000, 1000, 500, 100 | 0.0109 | 0.0129 | -0.002 |
| i/p, 2000, 1000, 500, 100 | 0.0105 | 0.0117 | -0.0013 |
| i/p, 1500, 500, 100 | 0.0095 | 0.0118 | -0.0023 |
| i/p, 1000, 100 | 0.0114 | 0.0124 | -0.001 |
| i/p, 4000, 2000, 1000, 500 | 0.0098 | 0.0126 | -0.0027 |
| i/p, 3000, 1500, 1000, 500 | 0.0101 | 0.0124 | -0.0023 |
| i/p, 2000, 1000, 500 | 0.0091 | 0.0116 | -0.0025 |

| | | | |
|---|---|---|---|
| i/p, 1500, 1000, 500 | 0.0094 | 0.0119 | -0.0025 |
| i/p, 2000, 500 | 0.0094 | 0.012 | -0.0026 |
| i/p, 4000, 2000, 1000 | 0.0077 | 0.0127 | -0.0051 |
| i/p, 3000, 1500, 1000 | 0.0085 | 0.012 | -0.0035 |
| i/p, 2000, 1000 | 0.0087 | 0.0119 | -0.0032 |
| i/p, 1500, 1000 | 0.0089 | 0.0113 | -0.0025 |

**Supplementary Table S4**: Comparing the clustering scores between the clusters ($K$ = 13) obtained using iClusterPlus and the proposed technique applied on the multi-omics data.

| | **Silhouette width** | **Calinski-Harabasz index** |
|---|---|---|
| iCluster | 0.06 | 414.03 |
| Proposed technique | 0.24 | 565.31 |

**Supplementary Table S5**: Number of differentially expressed features in each datatype (PcG: protein-coding genes, lnc RNA: long non-coding RNA, miRNA: micro-RNA).

| **Cluster** | **PcG** | **lnc RNA** | **miRNA** | **DNA methylation** | **Protein expression** |
|---|---|---|---|---|---|
| C1 (Brain cluster) | 734 | 93 | 9 | 640 | 65 |
| C2 (GYN cluster) | 172 | 23 | 3 | 284 | 0 |
| C3 (Thyroid cluster) | 175 | 36 | 6 | 262 | 32 |
| C4 (Kidney cluster) | 216 | 39 | 4 | 134 | 22 |
| C5 (Breast cluster) | 149 | 21 | 3 | 189 | 5 |
| C6 (Prostate cluster) | 170 | 43 | 4 | 372 | 20 |
| C7 (GI cluster) | 202 | 37 | 8 | 1240 | 3 |
| C8 (Bladder cluster) | 156 | 33 | 9 | 931 | 5 |
| C9 (Squamous cluster) | 221 | 30 | 6 | 157 | 4 |
| C10 (Low CNV) | 58 | 16 | 0 | 45 | 1 |
| C11 (M2 phenotype) | 193 | 27 | 48 | 11 | 12 |
| C12 (Liver cluster) | 285 | 25 | 4 | 591 | 46 |
| C13 (Mixed) | 137 | 19 | 0 | 16 | 1 |

**Supplementary Table S6**: Top five GO process in each ML cluster identified by metascape analysis.

| **Cluster** | **GO Biological Processes** |
|---|---|
| C1 (Brain cluster) | modulation of chemical synaptic transmission, synaptic signaling, regulation of membrane potential, neuron projection development, regulation of ion transport |
| C2 (GYN cluster) | microtubule bundle formation, pattern specification process, skeletal muscle contraction, muscle organ development, heart development |
| C3 (Thyroid cluster) | regulation of thyroid hormone generation, cell-cell adhesion, regulation of dopamine receptor signaling pathway, cyclic nucleotide catabolic process, formation of primary germ layer |
| C4 (Kidney cluster) | anion transport, circulatory system process, renal system process, anion transmembrane transport, xenobiotic detoxification |
| C5 (Breast cluster) | embryonic morphogenesis, skeletal system morphogenesis, regulation of biomineral tissue development, prostate gland development, urogenital system development |
| C6 (Prostate cluster) | metal ion transport, myofibril assembly, secretion, urogenital system development, muscle system process |
| C7 (GI cluster) | epithelial cell differentiation, regulation of microvillus organization, cytokine-mediated signaling pathway, gland development, circulatory system process |
| C8 (Bladder cluster) | epithelial cell differentiation, multicellular organismal water homeostasis, skin development, sodium ion homeostasis, tube morphogenesis |
| C9 (Squamous cluster) | epidermis development, cell-cell adhesion, establishment of skin barrier, desmosome organization, regulation of endopeptidase activity |
| C10 (Low CNV) | regulation of blood pressure, response to ketone, import into cell, cellular response to tumor necrosis factor, response to metal ion |
| C11 (M2 phenotype) | phenol-containing compound biosynthetic process, skeletal system development, sensory organ development, neuropeptide signaling pathway, behavior |
| C12 (Liver cluster) | monocarboxylic acid metabolic process, small molecule catabolic process, organic hydroxy compound metabolic process, olefinic compound metabolic process, lipid homeostasis |
| C13 (Mixed) | lymphocyte activation, adaptive immune response, regulation of leukocyte activation, regulation of immune effector process, alpha-beta T cell activation |

**Supplementary Table S7**: Positively enriched pathways in each subgroup obtained by GSEA analysis using hallmark geneset (FDR q <= 0.05).

| Cluster | Pathways |
|---|---|
| C5 (Breast cluster) | HALLMARK_ESTROGEN_RESPONSE_EARLY (q: 0.002) |
| C6 (Prostate cluster) | HALLMARK_ANDROGEN_RESPONSE (q: 0.000) |
| C9 (Squamous cluster) | HALLMARK_MTORC1_SIGNALING (q: 0.010),<br>HALLMARK_G2M_CHECKPOINT (q: 0.008),<br>HALLMARK_E2F_TARGETS (q: 0.007),<br>HALLMARK_MYC_TARGETS_V1 (q: 0.010),<br>HALLMARK_P53_PATHWAY (q: 0.040) |
| C12 (Liver cluster) | HALLMARK_XENOBIOTIC_METABOLISM (q: 0.000),<br>HALLMARK_BILE_ACID_METABOLISM (q: 0.000),<br>HALLMARK_FATTY_ACID_METABOLISM (q: 0.000),<br>HALLMARK_COAGULATION (q: 0.003) |
| C13 (Mixed) | HALLMARK_ALLOGRAFT_REJECTION (q: 0.000),<br>HALLMARK_IL6_JAK_STAT3_SIGNALING (q: 0.000),<br>HALLMARK_INFLAMMATORY_RESPONSE (q: 0.000),<br>HALLMARK_INTERFERON_GAMMA_RESPONSE (q: 0.000),<br>HALLMARK_IL2_STAT5_SIGNALING (q: 0.000),<br>HALLMARK_COMPLEMENT (q: 0.001),<br>HALLMARK_TNFA_SIGNALING_VIA_NFKB (q: 0.005),<br>HALLMARK_INTERFERON_ALPHA_RESPONSE (q: 0.008),<br>HALLMARK_KRAS_SIGNALING_UP (q: 0.040) |

**Supplementary Table S8**: Negatively enriched pathways in each subgroup obtained by GSEA analysis using hallmark geneset (FDR q <= 0.05).

| Cluster | Pathways |
|---|---|
| C2 (GYN cluster) | HALLMARK_KRAS_SIGNALING_UP (q: 0.010) |
| C3 (Thyroid cluster) | HALLMARK_E2F_TARGETS (q: 0.030),<br>HALLMARK_G2M_CHECKPOINT (q: 0.010),<br>HALLMARK_MTORC1_SIGNALING (q: 0.010),<br>HALLMARK_MYC_TARGETS_V2 (q: 0.040),<br>HALLMARK_MITOTIC_SPINDLE (q: 0.050) |
| C4 (Kidney cluster) | HALLMARK_G2M_CHECKPOINT (q: 0.003),<br>HALLMARK_E2F_TARGETS (q: 0.006),<br>HALLMARK_MYC_TARGETS_V1 (q: 0.030),<br>HALLMARK_UNFOLDED_PROTEIN_RESPONSE (q: 0.040),<br>HALLMARK_MITOTIC_SPINDLE (q: 0.030) |
| C11 (M2 phenotype) | HALLMARK_ESTROGEN_RESPONSE_LATE (q: 0.010),<br>HALLMARK_ESTROGEN_RESPONSE_EARLY (q: 0.010) |

**Supplementary Table S9**: Immune cells from LM22 gene signature enriched in each subgroup identified by Cibersort analysis.

| **Cluster** | **Immune cell type** |
|---|---|
| C1 (Brain cluster) | Monocytes, Mast cells activated, Eosinophils |
| C4 (Kidney cluster) | T-cells gamma delta, T-cells CD8 |
| C6 (Prostate cluster) | T-cells CD4 memory resting, Mast cells resting |
| C7 (GI cluster) | T-cells CD4 memory activated, B-cells naive |
| C8 (Bladder cluster) | Dendritic cells activated, T-cells CD4 naive |
| C9 (Squamous cluster) | NK cells resting, T-cells CD4 memory activated |
| C10 (Low CNV) | B-cells memory, T-cells regulatory Tregs |
| C11 (M2 phenotype) | Macrophages M2 |
| C12 (Liver cluster) | NK cells activated |
| C13 (Mixed) | T-cells CD4 naive |

**Supplementary Fig. S1**: Steps followed in preprocessing each datatype.

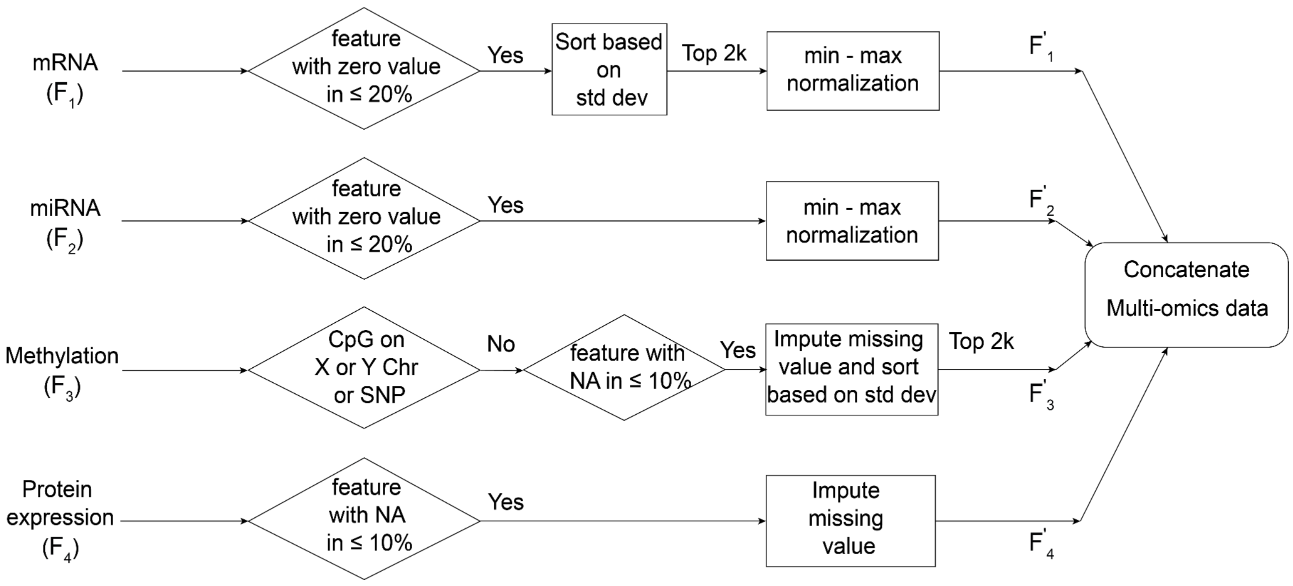


**Supplementary Fig. S2**: (a) Figures showing the CDF curves for consensus clustering from *K* = 2 to 20. KM curves for (b) overall survival (OS) and (c) disease-free survival (DFS) in the ML clusters.

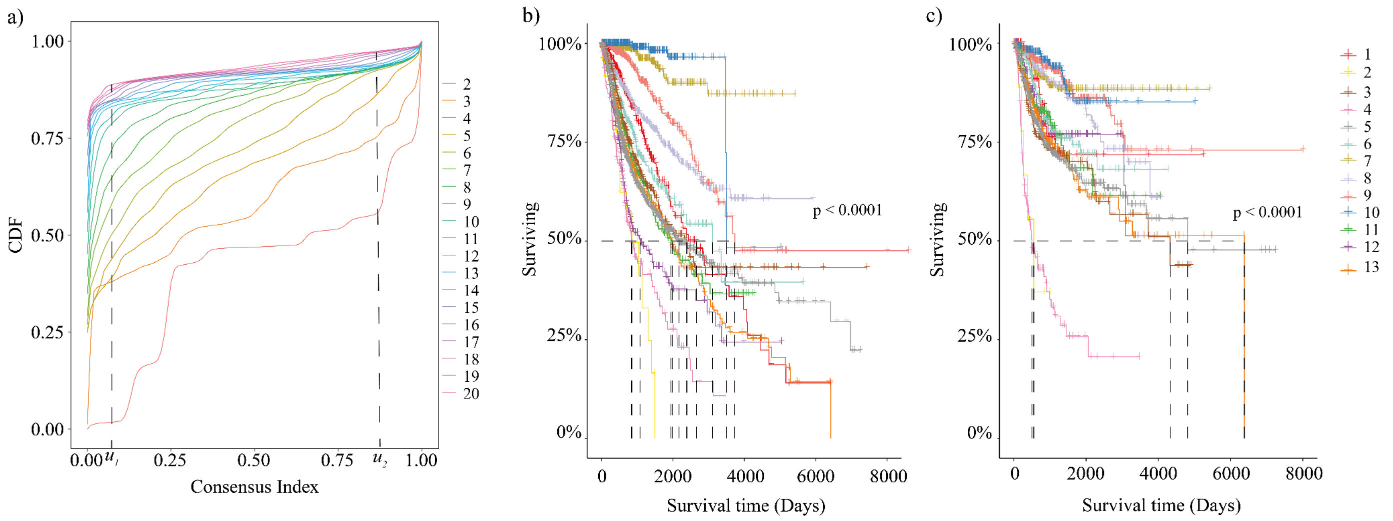

**Supplementary Fig. S3**: Figure showing the proportion of samples from each tumor type in each ML cluster.

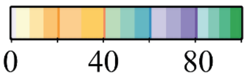


| | C1 (402) | C2 (372) | C3 (360) | C4 (489) | C5 (471) | C6 (342) | C7 (538) | C8 (250) | C9 (688) | C10 (26) | C11 (461) | C12 (169) | C13 (1135) |
|---|---|---|---|---|---|---|---|---|---|---|---|---|---|
| ACC (45) | 0 | 0 | 0 | 0 | 0 | 0 | 0 | 0 | 0 | 0 | 9.5 | 0 | 0.088 |
| BLCA (336) | 0 | 0.81 | 0 | 0 | 0 | 0 | 0.37 | 98 | 3.6 | 0 | 0 | 0 | 5.3 |
| BRCA (605) | 0 | 3.8 | 0 | 0 | 100 | 0 | 0 | 0 | 3.9 | 0 | 0 | 0 | 8.3 |
| CESC (160) | 0 | 2.2 | 0 | 0 | 0 | 0 | 3.2 | 0 | 18 | 0 | 0 | 0 | 0.97 |
| CHOL (30) | 0 | 0 | 0 | 0 | 0 | 0 | 1.5 | 0 | 0 | 23 | 0 | 5.3 | 0.62 |
| COAD (224) | 0 | 0 | 0 | 0 | 0 | 0 | 43 | 0 | 0 | 0 | 0 | 0 | 0.97 |
| DLBC (33) | 0 | 0 | 0 | 0 | 0 | 0 | 0 | 0 | 2.6 | 3.8 | 0 | 0 | 1.2 |
| ESCA (105) | 0 | 0.54 | 0 | 0 | 0 | 0 | 6.1 | 0 | 7.7 | 0 | 0 | 0 | 1.5 |
| HNSC (323) | 0 | 0.54 | 0 | 0 | 0.21 | 0 | 0 | 0 | 40 | 0 | 0 | 0 | 4.2 |
| KICH (63) | 0 | 0 | 0 | 12 | 0 | 0 | 0 | 0 | 0 | 3.8 | 0.43 | 0 | 0 |
| KIRC (259) | 0 | 0 | 0 | 50 | 0 | 0 | 0 | 0 | 0 | 0 | 0.65 | 0 | 1.1 |
| KIRP (203) | 0 | 0 | 0 | 38 | 0 | 0 | 0 | 0.8 | 0 | 38 | 0 | 0 | 0.53 |
| LGG (418) | 100 | 0 | 0 | 0 | 0 | 0 | 0 | 0 | 0 | 0 | 3.3 | 0 | 0.088 |
| LIHC (175) | 0 | 0 | 0 | 0 | 0 | 0 | 0 | 0 | 0 | 15 | 0 | 95 | 0.97 |
| LUAD (313) | 0 | 0.54 | 0 | 0 | 0 | 0 | 1.7 | 0.8 | 0.58 | 3.8 | 0 | 0 | 26 |
| LUSC (252) | 0 | 1.1 | 0 | 0 | 0 | 0 | 0 | 0 | 22 | 3.8 | 0.22 | 0 | 8.1 |
| MESO (60) | 0 | 0 | 0 | 0 | 0 | 0 | 0 | 0 | 0 | 0 | 6.7 | 0 | 2.6 |
| PAAD (96) | 0 | 0 | 0 | 0 | 0 | 0 | 2.4 | 0 | 0 | 0 | 0.43 | 0 | 7.1 |
| PCPG (77) | 0 | 0 | 0 | 0 | 0 | 0 | 0 | 0 | 0 | 0 | 16 | 0 | 0.088 |
| PRAD (343) | 0 | 0 | 0 | 0 | 0 | 100 | 0 | 0 | 0 | 0 | 0 | 0 | 0.088 |
| READ (75) | 0 | 0 | 0 | 0 | 0 | 0 | 14 | 0 | 0 | 0 | 0 | 0 | 0 |
| SARC (216) | 0 | 0.27 | 0 | 0 | 0 | 0 | 0 | 0 | 0.15 | 3.8 | 39 | 0 | 2.7 |
| SKCM (85) | 0 | 0 | 0 | 0 | 0 | 0 | 0 | 0 | 0.15 | 3.8 | 16 | 0 | 0.79 |
| STAD (261) | 0 | 0.81 | 0 | 0 | 0 | 0 | 27 | 0 | 0.29 | 0 | 0 | 0 | 9.6 |
| TGCT (118) | 0 | 0.27 | 0 | 0 | 0 | 0 | 0 | 0 | 0 | 0 | 4.1 | 0 | 8.6 |
| THCA (365) | 0 | 0 | 100 | 0 | 0 | 0 | 0 | 0 | 0 | 0 | 0 | 0 | 0.44 |
| THYM (86) | 0 | 0 | 0 | 0 | 0 | 0 | 0 | 0 | 0.87 | 0 | 0 | 0 | 7 |
| UCEC (298) | 0 | 77 | 0 | 0 | 0 | 0 | 0.19 | 0 | 0 | 0 | 0 | 0 | 0.79 |
| UCS (47) | 0 | 12 | 0 | 0 | 0 | 0 | 0 | 0 | 0.15 | 0 | 0 | 0 | 0.18 |
| UVM (12) | 0 | 0 | 0 | 0 | 0 | 0 | 0 | 0 | 0 | 0 | 2.6 | 0 | 0 |

**Supplementary Fig. S4**: Steps followed in preprocessing each datatype for molecular characterization.

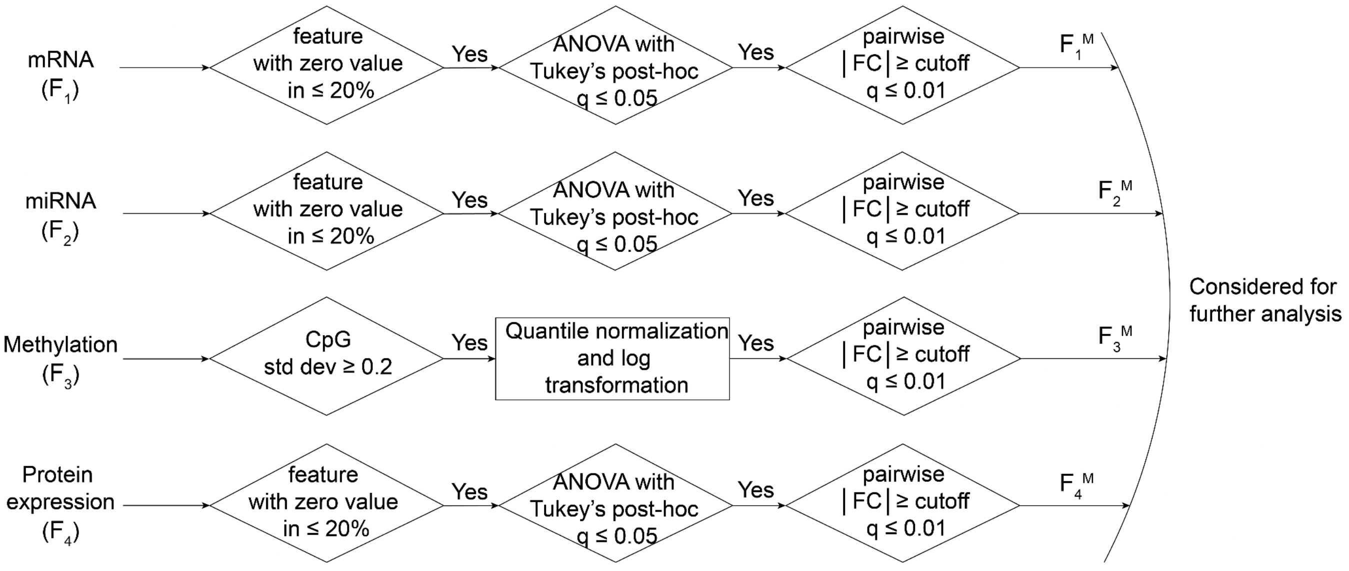


**Supplementary Fig. S5**: Heatmap showing the distribution of LM22 immune cells in each cluster.

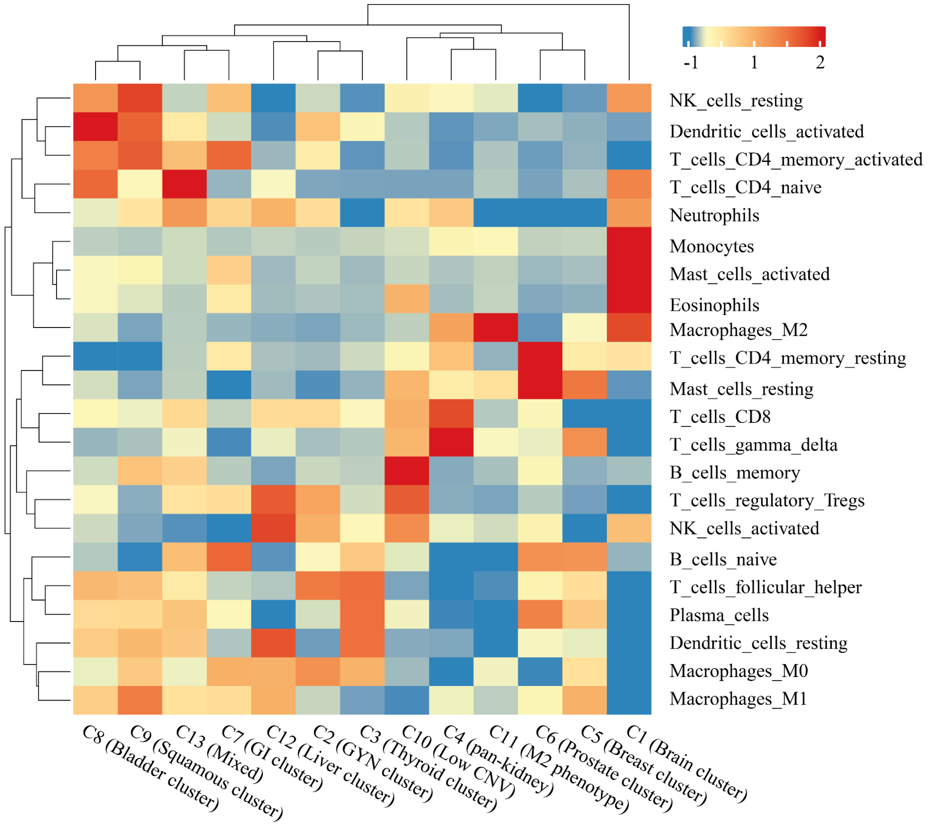

**Supplementary Fig. S6:** G-score of top 5 segments in each ML cluster indicating the copy number aberrations. (a) to (m): ML cluster 1 to ML cluster 13.

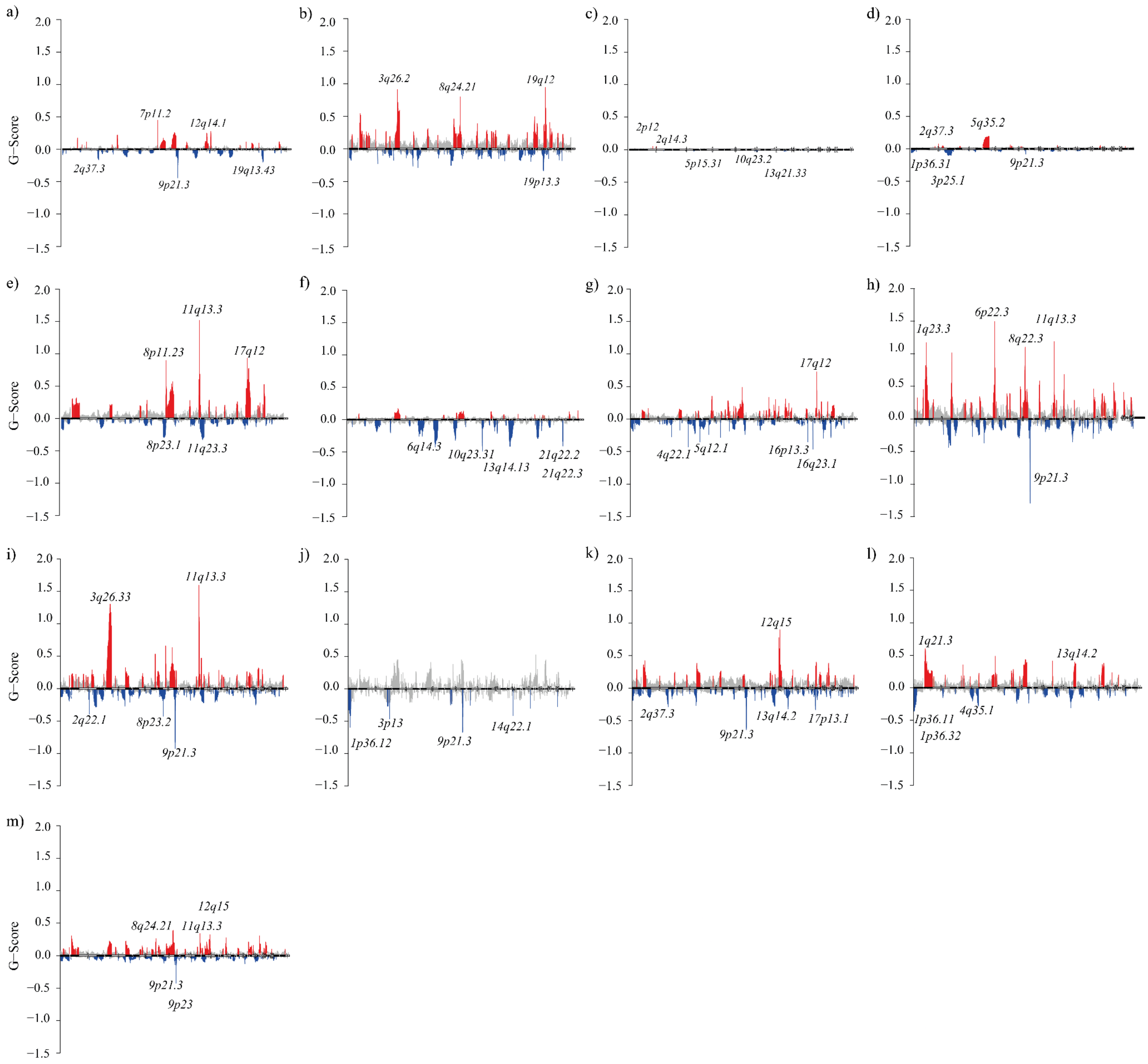

**Supplementary Fig. S7**: Pathway analysis of ML clusters. (a) Heatmap showing enrichment of different gene programs in all clusters. (b) Heatmap showing enrichment of different drug programs in all clusters.

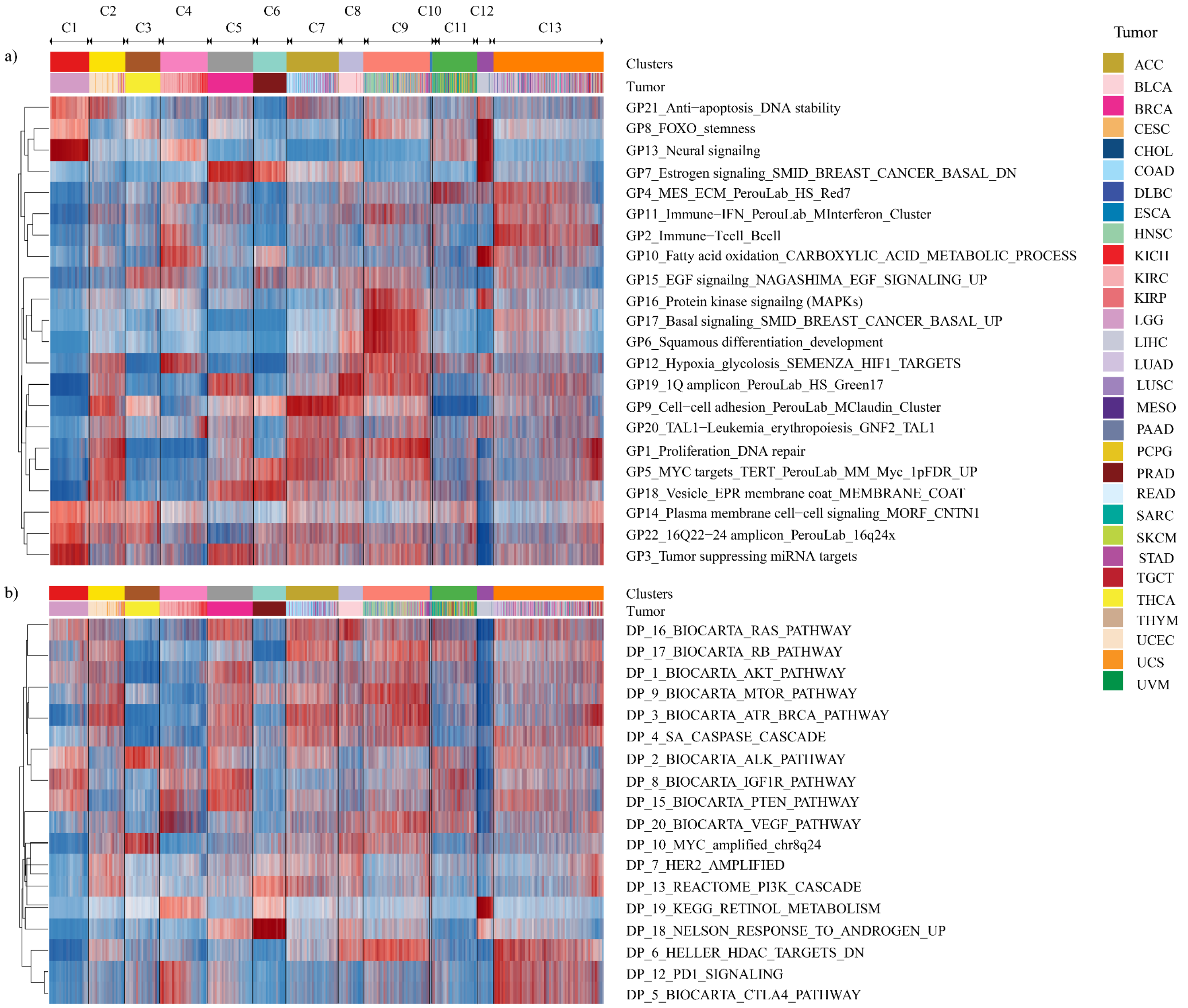

**Supplementary Fig. S8**: Figure showing the proportion of samples from each tumor type in each ML cluster in the validation data set.

Colour scale: 0 40 80

| | C1(210) | C2(169) | C3(140) | C4(131) | C5(138) | C6(153) | C7(225) | C8(59) | C9(399) | C10(2) | C11(311) | C12(198) | C13(523) |
|---|---|---|---|---|---|---|---|---|---|---|---|---|---|
| ACC (35) | 0 | 0 | 0 | 0 | 0 | 0 | 0 | 0 | 0 | 0 | 11 | 0 | 0.19 |
| BLCA (76) | 0 | 1.2 | 0 | 0 | 0 | 0 | 0.44 | 97 | 0.75 | 0 | 0.32 | 0 | 2.3 |
| BRCA (177) | 0 | 2.4 | 0 | 0 | 100 | 0 | 0 | 0 | 2.5 | 0 | 0.96 | 0 | 4.2 |
| CESC (147) | 0 | 4.7 | 0 | 0 | 0 | 0 | 9.3 | 0 | 26 | 0 | 0 | 0 | 2.7 |
| CHOL (6) | 0 | 0 | 0 | 0 | 0 | 0 | 0.44 | 0 | 0 | 50 | 0 | 0.51 | 0.57 |
| COAD (52) | 0 | 0 | 0 | 0 | 0 | 0 | 22 | 0 | 0 | 0 | 0 | 0 | 0.38 |
| DLBC (15) | 0 | 0 | 0 | 0 | 0 | 0 | 0 | 0 | 2.5 | 0 | 0 | 0 | 0.96 |
| ESCA (80) | 0 | 0 | 0 | 0 | 0 | 0 | 15 | 0 | 7.3 | 0 | 0 | 0 | 3.4 |
| GBM (140) | 54 | 0 | 0 | 0 | 0 | 0 | 0 | 0 | 0 | 0 | 6.8 | 0 | 1.1 |
| HNSC (205) | 0 | 0.59 | 0 | 0 | 0 | 0 | 0 | 1.7 | 43 | 0 | 0 | 0 | 6.3 |
| KICH (3) | 0 | 0 | 0 | 2.3 | 0 | 0 | 0 | 0 | 0 | 0 | 0 | 0 | 0 |
| KIRC (60) | 0 | 0 | 0 | 44 | 0 | 0 | 0 | 0 | 0 | 0 | 0.32 | 0 | 0.38 |
| KIRP (72) | 0 | 0 | 0 | 54 | 0 | 0 | 0 | 1.7 | 0 | 0 | 0 | 0 | 0 |
| LGG (98) | 46 | 0 | 0 | 0 | 0 | 0 | 0 | 0 | 0 | 0 | 0.32 | 0 | 0 |
| LIHC (202) | 0 | 0 | 0 | 0 | 0 | 0 | 0 | 0 | 0 | 50 | 0 | 99 | 0.76 |
| LUAD (145) | 0 | 0 | 0 | 0 | 0 | 0 | 0 | 0 | 0 | 0 | 0 | 0 | 28 |
| LUSC (118) | 0 | 1.8 | 0 | 0 | 0 | 0 | 0 | 0 | 18 | 0 | 0 | 0 | 8.2 |
| MESO (27) | 0 | 0 | 0 | 0 | 0 | 0 | 0 | 0 | 0 | 0 | 3.5 | 0 | 3.1 |
| OV (10) | 0 | 5.9 | 0 | 0 | 0 | 0 | 0 | 0 | 0 | 0 | 0 | 0 | 0 |
| PAAD (88) | 0 | 0 | 0 | 0 | 0 | 0 | 6.2 | 0 | 0 | 0 | 1.9 | 0 | 13 |
| PCPG (102) | 0 | 0 | 0 | 0 | 0 | 0 | 0 | 0 | 0 | 0 | 33 | 0 | 0 |
| PRAD (155) | 0 | 0 | 0 | 0 | 0 | 100 | 0 | 0 | 0 | 0 | 0 | 0 | 0.38 |
| READ (23) | 0 | 0 | 0 | 0 | 0 | 0 | 10 | 0 | 0 | 0 | 0 | 0 | 0 |
| SARC (45) | 0 | 0.59 | 0 | 0 | 0 | 0 | 0 | 0 | 0 | 0 | 13 | 0 | 0.76 |
| SKCM (20) | 0 | 0 | 0 | 0 | 0 | 0 | 0 | 0 | 0 | 0 | 6.1 | 0 | 0.19 |
| STAD (134) | 0 | 0.59 | 0 | 0 | 0 | 0 | 36 | 0 | 0 | 0 | 0 | 0 | 9.8 |
| TGCT (32) | 0 | 0 | 0 | 0 | 0 | 0 | 0 | 0 | 0 | 0 | 1.3 | 0 | 5.4 |
| THCA (142) | 0 | 0 | 100 | 0 | 0 | 0 | 0 | 0 | 0 | 0 | 0 | 0 | 0.38 |
| THYM (38) | 0 | 0 | 0 | 0 | 0 | 0 | 0 | 0 | 0.25 | 0 | 0 | 0 | 7.1 |
| UCEC (133) | 0 | 76 | 0 | 0 | 0 | 0 | 0 | 0 | 0 | 0 | 0.32 | 0 | 0.57 |
| UCS (10) | 0 | 5.9 | 0 | 0 | 0 | 0 | 0 | 0 | 0 | 0 | 0 | 0 | 0 |
| UVM (68) | 0 | 0 | 0 | 0 | 0 | 0 | 0 | 0 | 0 | 0 | 22 | 0 | 0.19 |

**Supplementary Fig. S9**: Figure showing the activity of various (a) gene and (b) drug programs in the validation dataset.

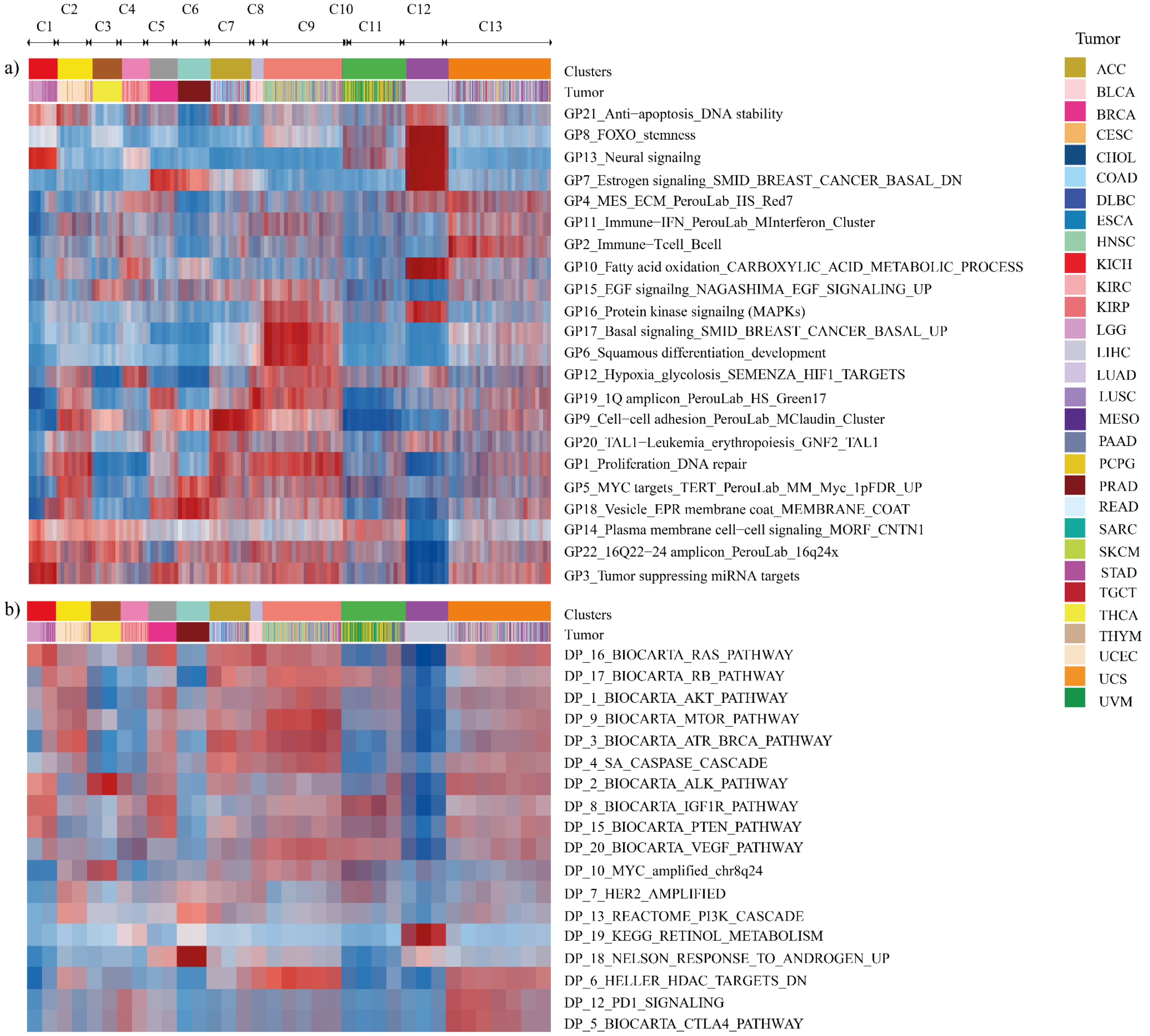

**The algorithm used to compute the values of α, β, and γ used to compute the prediction probabilities for the linear decision-level fused classification model.**

**Supplementary Algorithm S1:** Calculate α, β, and γ

$\alpha = 0: 0.01: 1$

**for** $\alpha_i$ in α:

  $\beta = 0 : 0.01 : (1 - \alpha_i)$

  **for** $\beta_j$ in β:

    $\gamma = 1 - \alpha_i - \beta_j$

  **end for**

**end for**
